# Biceps femoris long head sarcomere and fascicle length adaptations after three weeks of eccentric exercise training

**DOI:** 10.1101/2021.01.18.427202

**Authors:** Patricio A. Pincheira, Melissa A. Boswell, Martino V. Franchi, Scott L. Delp, Glen A. Lichtwark

## Abstract

**Purpose:** Eccentric exercise is widely used to increase muscle fascicle lengths and thus decrease the risk of muscle strain injuries. However, the mechanisms behind this protection are still unknown. The aim of this study was to determine whether Biceps femoris long head (BFlh) fascicle length increases in response to three weeks of eccentric exercise training are the result of addition of in-series sarcomeres within muscle fibres.

**Methods:** Ten recreationally active participants (age: 27 ± 3 years, mass: 70 ± 14 kg, height: 174 ± 9 cm) completed three weeks of Nordic hamstring exercise (NHE) training. We collected in vivo sarcomere and muscle fascicle images of the BFlh in two regions (central and distal), utilising microendoscopy and 3D ultrasonography. These images allowed us to estimate sarcomere length, sarcomere number and fascicle lengths before and after the training intervention.

**Results:** Eccentric knee flexion strength increased after the training (15%, P < 0.001, η_p_^2^= 0.75). Further, we found a significant increase in fascicle (21%, P < 0.001, η_p_^2^ = 0.81) and sarcomere (17%, P < 0.001, η_p_^2^ = 0.9) lengths in the distal but not in the central portion of the muscle. The estimated number of in series sarcomeres did not change in either region.

**Conclusion:** Fascicle length adaptations appear to be heterogeneous in the BFlf in response to three weeks of NHE training. An increase in sarcomere length, rather than the addition of sarcomeres in series, appears to be underlying this adaptation. The mechanism driving regional increases in fascicle and sarcomere length remain unknown, but we speculate it may be driven by regional changes in the passive tension of muscle or connective tissue adaptations.

## 1 Introduction

Eccentric training is beneficial in enhancing muscle strength, power and speed ^11^. It can improve muscle mechanical function to a greater extent than other exercise modalities (e.g. concentric and isometric) ^16^. Continued eccentric training can cause an adaptation in muscles that protects it from subsequent exercise-induced muscle damage ^23^. For instance, training programmes that sufficiently load the eccentric phase of a movement induce changes in muscle fascicle lengths that are effective in the prevention of strain related injuries ^5,6,12,21^. Although these eccentric contraction-induced muscle modifications have been extensively studied ^11,16,36^, the mechanisms underlying fascicle length changes and the role of muscle architectural adaptations in preventing injuries are still largely unknown.

Nordic hamstring exercise (NHE) are one such eccentric exercise that elicits substantial changes in fascicle length, protecting the hamstring muscles from subsequent muscle damage or injury. Nevertheless, the macroscopic and microscopic changes that contribute to the mechanical load-induced hamstrings lengthening response remains poorly understood. The main hypothesis put forward is that repetitive eccentric contractions cause addition of sarcomeres in series within muscle fibres ^34,35^, reducing the likelihood of overstretching the muscle in subsequent eccentric loading. However, this hypothesis has not been tested due to the lack of simultaneous in vivo measurements of fascicle and sarcomere changes in human subjects. Additionally, methodological limitations, such as extrapolation when measuring fascicle length ^15^, may hamper interpretation of previous findings ^41,44^. Establishing the mechanisms of fascicle length adaptations after eccentric exercise, including the potential role of sarcomerogenesis, is needed to provide a scientific basis for designing improved training techniques.

As muscle remodelling processes appear to be related to the amount of stress and strain applied onto the muscle ^24^, it is also possible that muscle strain may be heterogeneous across different regions of the same muscle ^14,25^ implying that morphological, structural and even molecular adaptations due to eccentric training are non-uniform within the muscle ^3,17^. For example, in vivo measures of sarcomere length in the human tibialis anterior muscle have shown that sarcomeres are longer in the distal compared to the proximal region, despite fibres being of similar length ^27^. This implies that the force generating capacity of different regions of a muscle may be different, and therefore, different stress and strain distributions could exist. Animal studies also suggest that different muscle architectures and fibre lengths may influence if and how serial sarcomere adaptations occur ^33^. This highlights the need to use multiscale measurements, from micro to macro-organisation, to understand the nature of region-specific adaptations due to eccentric exercise.

While the in-vivo measurement of sarcomere and fascicle length has historically been unattainable, promising new methods have emerged to examine both fascicle length and individual sarcomere number in live subjects. Free hand 3D ultrasound (3DUS) combines 2D ultrasound scanning and 3D motion analysis to provide direct in vivo measurements of fascicle length within a wide area of tissue structure ^2^. As such, it is possible to measure the lengths of fascicles that are much longer than the field of view of conventional B mode imaging ^15^. Minimally invasive optical microendoscopy visualizes in vivo individual sarcomeres and their dynamical length variations ^28^. Combining freehand 3D ultrasound with optical microendoscopy may provide direct evidence of whether eccentric exercise induces adaptations in sarcomere number (sarcomerogenesis) in live human muscle.

The aim of this study was to measure multi-scale biceps femoris long head (BFlh) adaptations in response to NHE. We provide the first simultaneous in vivo measurement of both fascicle length and sarcomere length in multiple human BFlh muscle regions before and after an eccentric training intervention. We sought to investigate adaptions in the hamstring muscle because of the large increases in fascicle length that have previously been reported to occur after relatively short intervention periods (e.g. two weeks) ^41^. Also, the hamstrings is an important muscle group to study, considering that BFlh strains are the most prevalent non-contact injury in sports involving running ^13,38^. We hypothesised that in BFlh, fascicle length would increase in response to NHE and that such adaptations will be accompanied by an addition of sarcomeres in series within fascicles. A better understanding of how different regions of human muscle adapt to unaccustomed eccentric exercise may help to better understand strain injury mechanisms and to develop better injury preventing strategies.

## 2 Methods

### 2.1 Participants

Ten recreationally active volunteers (three female, age 25 ± 1 years, mass 55 ± 3 kg, height 164 ± 4 cm; seven male, age 29 ± 4 years, mass 77 ± 11 kg, height 181 ± 9 cm) participated in the study and were recruited between June and July of 2019. Participants were excluded if they had any lower limb, trunk, or wrist injuries within the past 18 months or performed NHE within the past 6 months. Participants provided written, informed consent to participate, but they were not involved in the study design or interpretation. The protocol was approved by the University of Queensland Institutional Review Board and Stanford Human Research Ethics Committee and was conducted according to the Declaration of Helsinki.

### 2.2 Study design

The NHE training lasted three weeks, a period previously reported to be sufficient to elicit lengthening of passively measured BFlh fascicles ^41^. On Day 1, baseline measurements of BFlh fascicle and sarcomere lengths were conducted. Then, participants were familiarised with the NHE device and underwent a baseline assessment of eccentric knee flexor strength. All measurements were performed on the right leg, the dominant leg for all participants (i.e. the leg that participants used to kick a ball). On Day 2, fascicle length was measured again, prior to the first NHE training session. Participants underwent a total of nine training sessions in the following three weeks (Table 1). Post-training BFlh fascicle and sarcomere lengths were measured one day after the last training session.

**Table 1.** Nordic hamstring exercise training intervention variables

| Week | Sessions per week | Sets | Repetitions | Total number of repetitions | Rest between sets |
| --- | --- | --- | --- | --- | --- |
| 1 | 3 | 4 | 6 | 72 | 2 min |
| 2 | 3 | 4 | 6 | 72 | 2 min |
| 3 | 3 | 4 | 8 | 96 | 2 min |

### 2.3 Nordic hamstring exercise training

Participants performed the NHE training using a custom made device similar to previous studies ^37,40,41^. Participants knelt on a padded board with their ankles secured with braces located superior to the lateral malleolus. We attached the ankle braces to uniaxial load cells (HT Sensor Technology, Shaanxi, RPC) secured to the board via a rod and swivel. From this position, participants slowly leaned forward with control to a metronome (∼ 30 degrees per second). To minimize concentric loading, once participants’ hands touched the floor, they pushed themselves back to the starting position with their arms and trunk. We provided strong verbal encouragement to participants during all testing and training sessions to ensure maximal effort in each repetition.

### 2.4 Eccentric knee flexor strength measurement

We measured hamstrings eccentric strength on the same custom device used for the training. Prior to testing, participants completed a warm-up set of four repetitions at 75% of their perceived maximum effort. After a three-minute rest period, participants then completed one set of four maximal NHE repetitions. Peak knee flexion moment was computed from the load cell measurement at the ankle. We analysed the average of the top three strength measurements of the four trials.

### 2.5 Fascicle length

We measured fascicle lengths using freehand 3D ultrasonography (3DUS) described elsewhere ^2,22^. The 3DUS scan involved B-mode ultrasound imaging (ArtUS EXT-1H + L15-7H40-A5 40mm transducer, Telemed, Vinilius, Lithuania) with synchronous optical tracking (Optitrack, Natural Point, OR, USA) of position and orientation of the transducer. The scan was collected and analysed using open-source software (Stradwin software v5.4, Mechanical Engineering, Cambridge University, UK). Before testing, we calibrated the system temporally and spatially as previously described ^2^. The 3DUS system has previously been reported to have excellent test-retest reliability for measuring muscle fascicle length ^18,22^.

During the 3DUS scans, participants laid prone with their hips and knees fully extended and their feet in a neutral position off the end of the bed. We quantitatively determined the BFlh scanning regions (Figure 1A). We first marked the position halfway between the ischial tuberosity and the top of the patella followed by distances 10% proximal and 30% distal to the midway point. We drew a line for the path of the BFlh between the proximal and distal ends of this scanning region. At a constant speed and pressure, we moved the ultrasound transducer over the scanning region in a transverse orientation that allowed identification of the collagenous tissue running along the length of the muscle. We performed six freehand 3DUS scans in each session. From each session, we selected the two scans with the best image quality (e.g., clear and visible fascicles and aponeuroses) for 2D image extraction.

**Figure 1.**
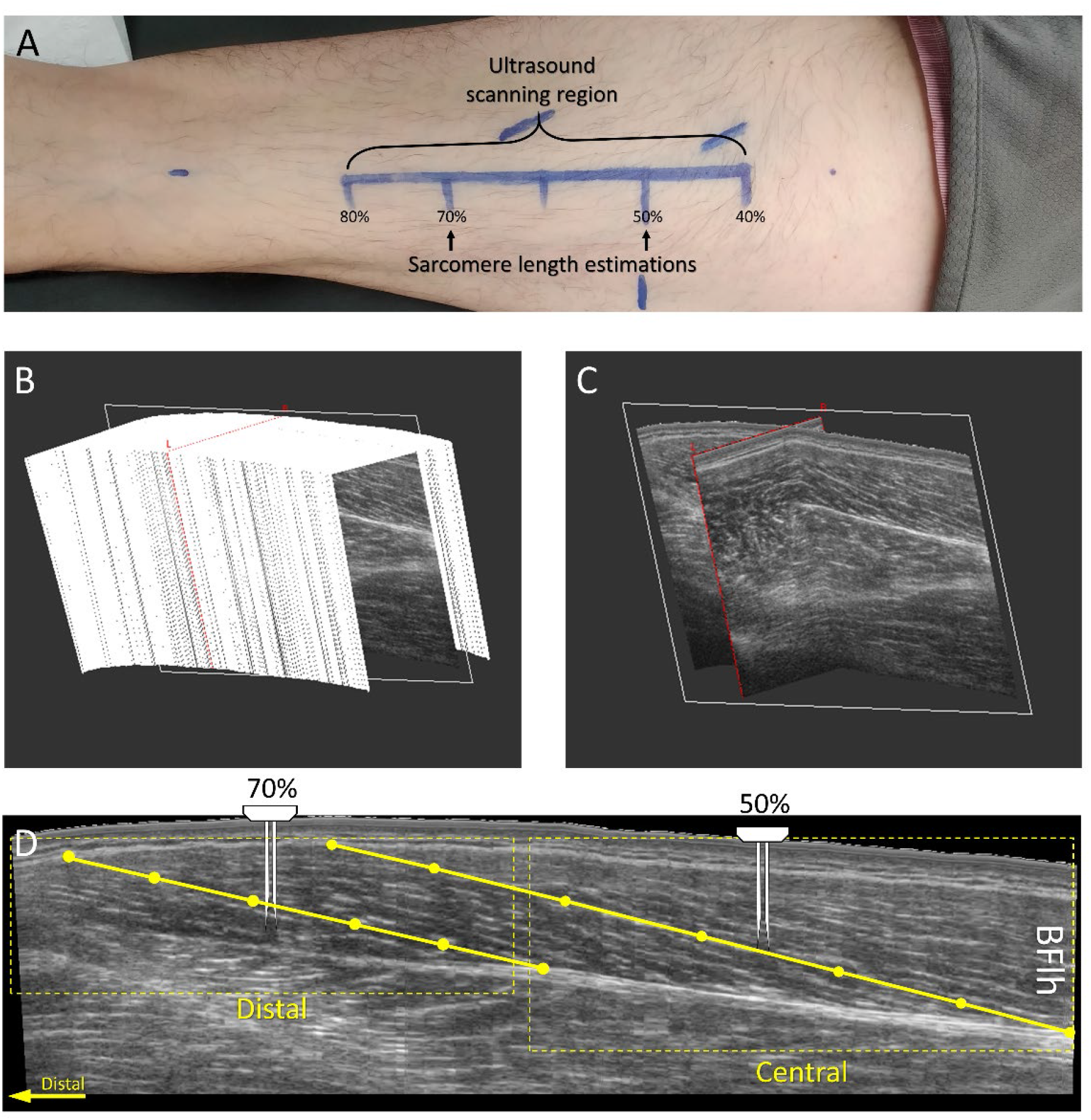
Experimental set-up for biceps femoris long head (BFlh) fascicle and sarcomere measurement. A) The ultrasound scanning region defined between 40% and 80% of the distance measured from the ischial tuberosity to the top of the patella. Sarcomere length estimations were made at 50 and 70% of the same distance. B) A stack of 2D B-mode ultrasound images from the BFlh from 3D ultrasonography. C) A 2D plane extracted from the stack of images. D) An image of the BFlh extracted from the 2D plane with the distal and central regions indicated by the dotted box. A muscle fascicle is shown by the yellow solid line, which we defined with 6 points. The needle probe for sarcomere imaging is shown at the locations of insertion.

From the stack of 2D B-mode ultrasound images collected (slice thickness ∼ 0.36 mm, Figure 1B), we selected a 2D plane that covered the full length of the BFlh fascicles (Figure 1C) and positioned it on the thickest part of the proximal cross-sectional area of the muscle. From this image, we determined the central and distal regions for fascicle length estimation (Figure 1D). Since fascicle orientation may change between regions of interest ^15,39^, we adjusted the location of the 2D plane separately for each region. In each image and region, we visually digitalized three entire fascicles using the 3D coordinates of six points per fascicle. We then imported the coordinates to Matlab (version 2015b, The Mathworks, MA, USA) and calculated fascicle length as the length of a parametric spline approximation fitted to the 3D coordinates. This method allowed us to consider fascicle curvature in our length estimations. Finally, we averaged the six fascicle length estimations in each region.

One researcher (PAP) completed all fascicle length measurements while blinded to participant ID and time. The test–retest reliability of fascicle length calculation between baseline and day 2 (Intra-class correlation ICC (3,k)), was “good” ^26^ (ICC = 0.85 confidence interval = 0.38 – 0.96).

### 2.6 Sarcomere length and number

We measured sarcomere lengths using second harmonic generation (SHG) microendoscopy (Zebrascope, Zebra Med Tech, CA) ^28,42^. The needle probe consisted of a pair of 2 cm needles with side-mounted lenses and 1 mm separation. One needle emits the laser light, while the other receives the SHG signals from the muscle fibres. The signal appears as repeating dark (myosin) and light bands. To reduce participant discomfort, we inserted the needle probe using a rapid, spring-loaded device. We defined two insertion sites: the central insertion site at 50% of the distance between the ischial tuberosity and the top of the patella, and the distal insertion site at 70% of the same distance from the ischial tuberosity (Figure 1A and 1D). The needle insertion sites were consistent with the central region of the fascicle length estimation sites. To ensure the fibres remained uncompromised, we inserted the probe such that the needles were approximately perpendicular to the fascicle plane (determined by ultrasound).

We analysed sarcomere lengths as described by Lichtwark et al. ^27^. We collected a sequence of images (image depth: 5–15 mm; sampling rate: 1 Hz) in real-time while slowly moving the microscope and needle through the muscle. During post-processing, we used a Gaussian filter to subtract white noise and a Fourier transform to calculate the strongest frequency spectrum between 1.5 and 5 μm. We selected a region on the sharpest fibre image for analysis. Across that region (∼100 μm in length), we repeated a Fourier transform to estimate the average sarcomere length. We took one measurement per individual fibre, totalling between 6 and 22 separate muscle fibre estimations (mean: 9.2 estimations) in each region per imaging session. Since the number of sarcomere length estimations varied between subjects, we used the median sarcomere length for each participant and each session for subsequent statistical analysis.

We estimated the number of sarcomeres in series for each fibre by dividing the fascicle length (used as a proxy for fibre length) by the median sarcomere length. One researcher (MAB) completed all sarcomere length calculations while blinded to participant ID and time.

### 2.7 Statistical analysis

We used a paired t-test to evaluate changes in eccentric knee flexor strength, fascicle length, sarcomere length, and the number of sarcomeres in series from baseline to the end of training. All statistical tests were conducted in Prism (version 8.4, GraphPad Software, CA, USA) with an alpha level set at P < 0.05. Effect sizes are presented as partial eta-squared (η_p_^2^). All values are presented as mean (standard deviation).

## 3 Results

Eccentric knee flexion strength was greater at the end of the three-week NHE training than at baseline (Baseline: 327.3 (85) Nm/Kg, End: 376 (67) Nm/Kg; P < 0.001, η_p_^2^ = 0.75). Fascicle length increased in the distal region (Baseline: 7.0 (0.9) cm, End: 8.5 (0.8) cm; P < 0.001, η_p_^2^ = 0.81), but not in the central region of the BFlh muscle (Baseline: 9.9 (0.7) cm, End: 10.1 (0.9) cm; P = 0.21, η_p_^2^ = 0.07) (Figure 2A). Similarly, sarcomere length increased in the distal region (Baseline: 2.9 (0.28) μm, End: 3.4 (0.23) μm; P < 0.001, η_p_^2^ = 0.9) but not in the central region (Baseline: 3.1 (0.17) μm, End: 3.2 (0.27) μm; P = 0.28, η_p_^2^ = 0.03) (Figure 2B). The estimated number of sarcomeres in series did not change in the distal region (Baseline: 24611 (4461) sarcomeres, End: 25382 (2799) sarcomeres; P = 0.22, η_p_^2^ = 0.07), nor the central region (Baseline: 32212 (2778) sarcomeres, End: 32092 (4271) sarcomeres; P = 0.46, η_p_^2^ < 0.01) (Figure 2C). In Figure 3, we present images displaying architectural adaptations for a representative subject. In Figure 4, we present all participants’ sarcomere length measurements, for all individual measures in each session and for each region of muscle.

**Figure 2.**
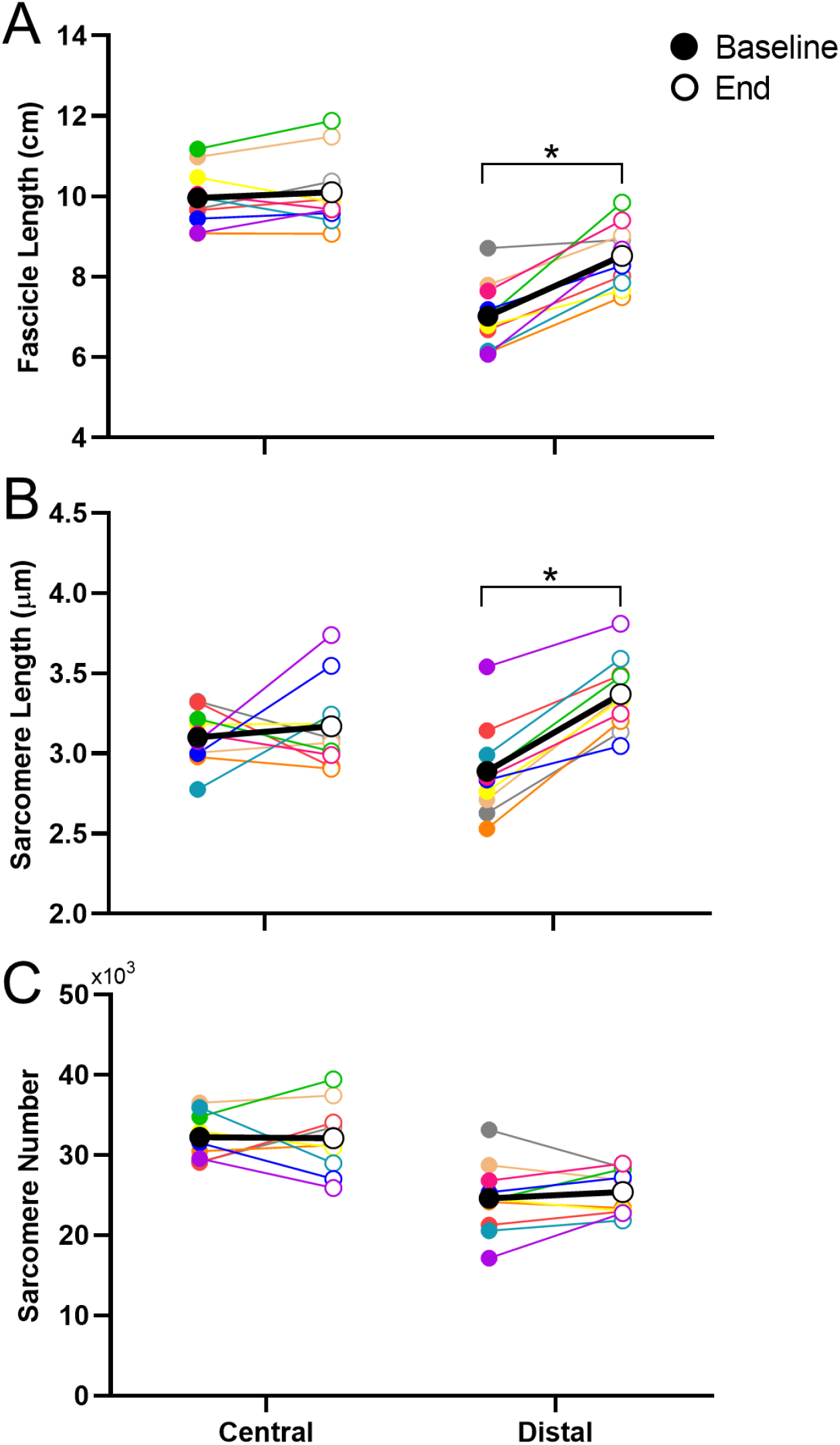
Biceps femoris long head architectural adaptations at group level. Fascicle lengths (A), sarcomere lengths (B) and sarcomere numbers (C) are presented for the central and distal portions of the muscle at baseline (filled circles) and after 9 sessions of Nordic hamstring exercise training (End, open circles). Each colour represents an individual participant. The average across all participant averages is presented in black. *Significant difference when comparing Baseline and End time points (P < 0.05).

**Figure 3.**
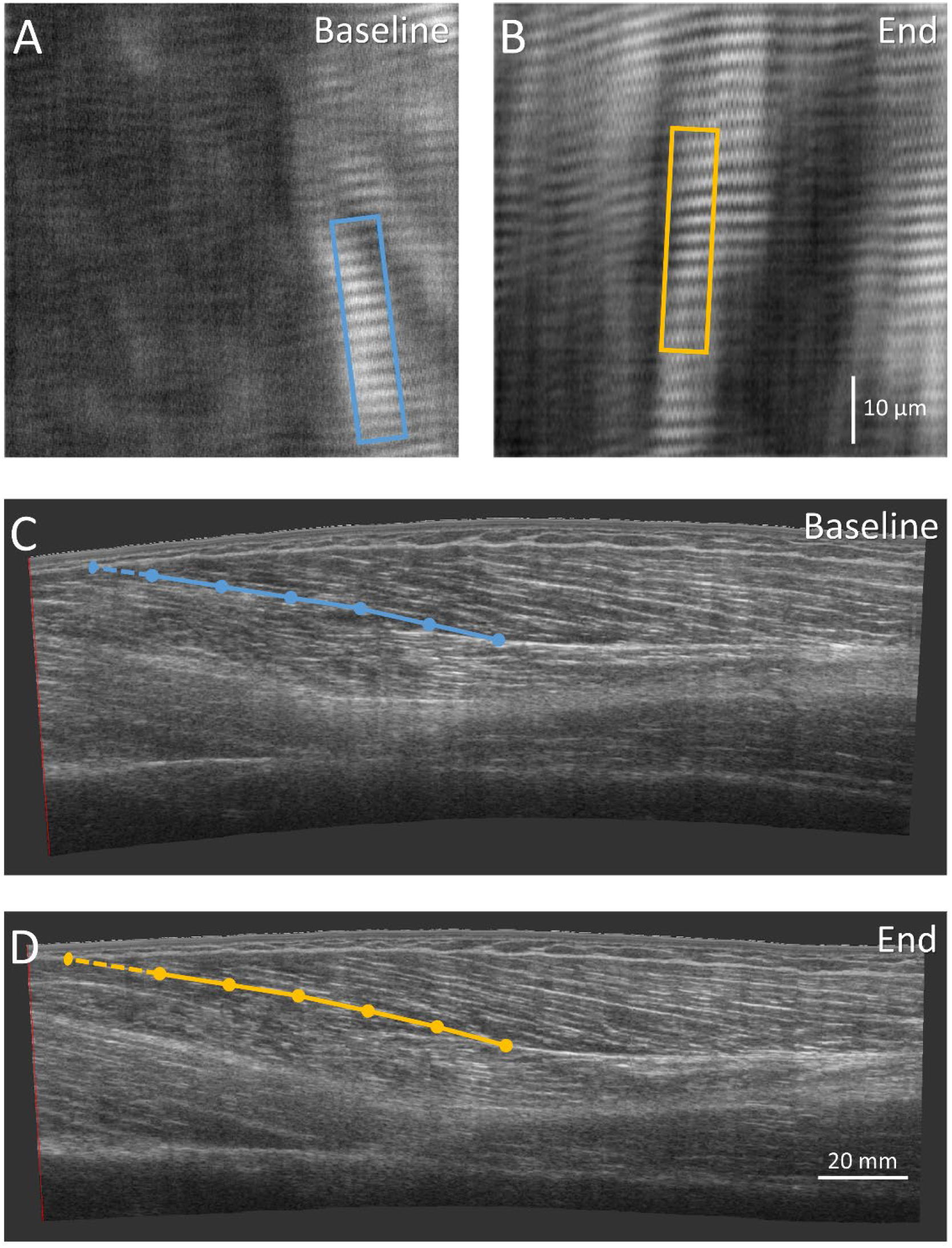
Images of the biceps femoris long head architectural adaptations for a representative subject. Sarcomere imaging at baseline (A) and at the end of the training (B) are presented for the distal portion of the muscle. The boxes represent the region where sarcomere length was estimated for each muscle fibre (Baseline, blue box: 2.9 μm; End, orange box: 3.6 μm). 2D sagittal-plane images at baseline (C) and at the end (D) created from 3D ultrasound are presented for the biceps femoris long head muscle. The overlayed lines represent a fascicle length estimation, with dots separating five segments of equal length. The last segment of the fascicle estimation line is drawn with a dotted line to highlight the change in fascicle length (Baseline: 93 mm; End: 101 mm).

**Fig 4.**
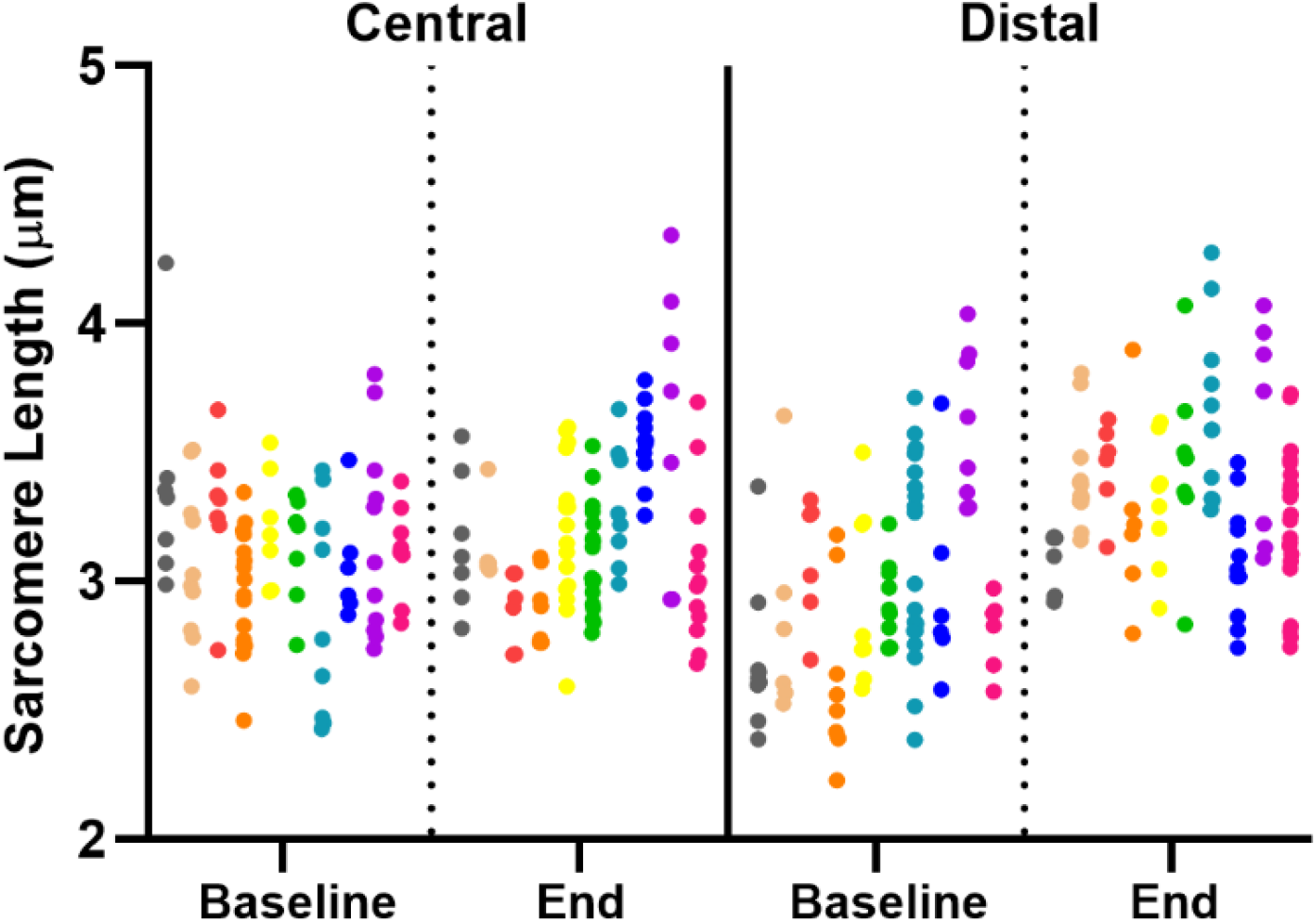
Sarcomere length measurements for baseline and the end of training for the central and distal regions. Individual measurements are represented by one dot and individual participants (N = 10) represented with different colours.

## 4 Discussion

In this study, we measured in vivo fascicle and sarcomere lengths in the human BFlh before and after an eccentric training intervention. Three weeks of NHE led to increased fascicle and sarcomere lengths in the distal portion of the BFlh. However, no changes in the number of sarcomeres in series was found. These findings provide new information about the early adaptations of the hamstring muscles in response to eccentric training.

### 4.1 Fascicle and sarcomere adaptations to eccentric training

Fascicle length in the BFlh increased after a short period of eccentric training (nine training sessions distributed over three weeks), similar to previous observations in the mid-portion of the muscle ^1,7,40,41,44^. However, we found heterogeneous fascicle length adaptations across the BFlh, as fascicle length increased in the distal but not in the central region of the muscle. The average change in fascicle length that we measured (∼ 1 cm) is smaller than the change observed previously (∼ 2 cm in the mid BFlh)^41^. These differences may be related to the imaging methods used, as previous studies ^1,7,40,41,44^ estimated muscle fascicle length using ultrasound imaging with a limited field of view of the muscle, which may overestimate changes in fascicle length ^15^.

We found that sarcomere length increased only in the distal portion of the muscle. In vivo measurement of both fascicle and sarcomere lengths allowed us to estimate the number of sarcomeres in series per fascicle. These estimates revealed no addition of sarcomeres in series in either region of muscle. Previous studies have suggested that sarcomerogenesis underlies the in-vivo changes in hamstrings muscle fascicle lengths after eccentric training ^1,7,41^. However, these assertions were supported indirectly, based on changes in fascicle lengths estimated with 2D ultrasonography ^1,7,41^ or changes in the angle-torque relationship ^9^. Thus, to our knowledge, this is the first study to show that eccentric training (such as three weeks of NHE) does not induce the addition of sarcomeres in series in humans, despite obvious site-specific increases in fascicle length. While this result may be unexpected, animal studies show that while some muscles experience sarcomerogenesis in response to eccentric exercise ^10,29^, others do not ^33^. Therefore, the results of this study call into question the predominate theory about how NHE might protect muscle from injury.

### 4.2 Interpretation of findings: Potential mechanisms of adaptation

While we did not see addition of serial sarcomeres in the muscle, the increase in fascicle lengths seen here and in previous studies ^1,7,40,41,44^ may represent adaptations to non-contractile tissue. Although the current study did not investigate non-contractile tissue adaptation, the change in length measured (in resting muscle) may represent changes in passive tension governed by adaptations in connective tissue, titin function, or both. Extracellular matrix components related to muscle fibre compliance (Tenascin-C), and force transmission to sarcomeres (Collagen XII), adapt quickly in response to eccentric exercise ^30^ and may account for our observations. Adaptations in titin stiffness or titin-actin interaction ^36^ seen after eccentric exercise, may also play a role in the increase in sarcomere and fascicle length. Furthermore, changes in tendon compliance ^20^ cannot be discarded as a source of adaptation. Overall, the mechanisms underpinning BFlh architectural adaptations are still not well understood and warrant further investigation.

The heterogeneous adaptation to fascicle length at different muscle regions that we observed may be important for understanding how a training stimulus relates to muscle adaptation. An heterogeneous distribution of fibre strain ^4^ and muscle activity ^19,31^ towards the distal portion of the BFlh may have influenced this region’s adaptations. Since hip position remains constant during the NHE, changes in fascicle excursion and tissue strain could be larger in the distal (shorter) portion of the BFlh muscle ^45^. Such differences in strain could provide stimulus (e.g. mechanotransduction ^17^) for differential adaptations in fascicle lengths across different muscle regions. Further studies are required to better understand the stress and strain distribution across the muscle during eccentric exercise, like the NHE, and how this relates to muscle adaptation.

### 4.3 Practical implications

Clinical interventions that employ Nordic hamstring exercise for three weeks can be expected to increase fascicle lengths only in the distal region of the BFlh. This finding presents a unique insight to the immediate (three weeks) adaptations that occur in response to NHE, which may be important for tailoring injury prevention training regimes, or identifying muscle at higher risk of injury. Previous studies suggests that increases in BFlh fascicle length may help protect athletes from hamstring injury ^12,21,43^; however, this study questions the proposed mechanism for such protection. While ‘long and strong’ ^8^ is considered an effective muscle adaptation to eccentric hamstring exercise, the goal of the exercises to induce these changes needs further consideration. While the addition of sarcomeres in series may play a role in longer-term training, we did not detect these changes early in the adaptive process. It is likely that there is little protection from injury offered to muscle due to sarcomerogenesis in this early stage of training, Analysis of NHE interventions of longer durations using sarcomere and fascicle imaging could provide essential evidence to further evaluate this commonly used training method.

### 4.4 Strengths and Limitations

Combining measurements in macro (fascicle) and micro (sarcomere) adaptations allowed us to overcome some of the methodological limitations of past studies, including assessment of the full length of the BFlh fascicles (considering fascicle curvature) in different muscle regions, and visualization of human in vivo sarcomere length adaptations. Nevertheless, we recognize several limitations of our study. First, this was a relatively short-term term training study; hence, we cannot rule out sarcomerogenesis in response to more prolonged training periods of various intensities. For instance, region-specific adaptations in mechanotransductor pathways related to muscle growth seems to peak at four and eight weeks after eccentric resistance training ^17^; thus, sarcomerogenesis may still occur at later stages in NHE training. As an additional limitation, we estimated only passive sarcomere and fascicle lengths ^32^. Future studies should consider in vivo assessment of sarcomere contractile dynamics ^42^ after eccentric training interventions.

## 5 Conclusion

NHE training (i.e. a total of 9 sessions in 3 weeks) led to an increase in fascicle and sarcomere lengths in the distal region of the BFlh muscle, but not the central region. These results suggest an heterogeneous distribution of strain and adaptation within the muscle. The addition of sarcomeres in series did not accompany the adaptations observed in either region. While the mechanisms behind these findings remain unclear, adaptations in connective tissue, titin, and extracellular matrix may play an important role.

